# Metagenomic sequencing reveals a lack of virus exchange between native and invasive freshwater fish across the Murray-Darling Basin, Australia

**DOI:** 10.1101/2021.02.25.432824

**Authors:** Vincenzo A. Costa, Jonathon C.O. Mifsud, Dean Gilligan, Jane E. Williamson, Edward C. Holmes, Jemma L. Geoghegan

## Abstract

Biological invasions are among the biggest threats to freshwater biodiversity. This is increasingly relevant in the Murray-Darling Basin, Australia, particularly since the introduction of the common carp (*Cyprinus carpio*). This invasive species now occupies up to 90% of fish biomass, with hugely detrimental impacts on native fauna and flora. To address the ongoing impacts of carp, *cyprinid herpesvirus 3* (*CyHV-3*) has been proposed as a potentially effective biological control. Crucially, however, it is unknown whether *CyHV-3* and other cyprinid herpesviruses already exist in the Murray-Darling. Further, little is known about those viruses that naturally occur in wild freshwater fauna, and the frequency with which these viruses jump species boundaries. To document the evolution and diversity of freshwater fish viromes and better understand the ecological context to the proposed introduction of *CyHV-3*, we performed a meta-transcriptomic viral survey of invasive and native fish across the Murray-Darling Basin, covering over 2,200 km of the river system. Across a total of 36 RNA libraries representing 10 species, we failed to detect *CyHV-3* nor any closely related viruses. Rather, meta-transcriptomic analysis identified 18 vertebrate-associated viruses that could be assigned to the *Arenaviridae, Astroviridae, Bornaviridae, Caliciviridae, Coronaviridae, Chuviridae, Flaviviridae, Hantaviridae, Hepeviridae, Paramyxoviridae, Picornaviridae, Poxviridae, Reoviridae* and *Rhabdoviridae* families, and a further 27 that were deemed to be associated with non-vertebrate hosts. Notably, we revealed a marked lack of viruses that are shared among invasive and native fishes sampled here, suggesting that there is little virus transmission from common carp to native fish species. Overall, this study provides the first data on the viruses naturally circulating in a major river system and supports the notion that fish harbour a large diversity of viruses with often deep evolutionary histories.

**Author Summary:** The ongoing invasion of the common carp in the Murray-Darling Basin, Australia, has wreaked havoc on native freshwater ecosystems. This has stimulated research into the possible biological control of invasive carp through the deliberate release of the virus *cyprinid herpesvirus 3* (*CyHV-3*). However, little is known on the diversity of viruses that naturally circulate in wild freshwater fauna, whether these viruses are transmitted between invasive and native species, nor if *CyHV-3* or other cyprinid herpesviruses are already present in the basin. To address these fundamental questions we employed meta-transcriptomic next-generation sequencing to characterise the total assemblage of viruses (i.e. the viromes) in three invasive and seven native fish species cohabiting at 10 sites across 2,200 km of the river system. From this analysis we identified 18 vertebrate-associated viruses across 14 viral families, yet a marked lack of virus transmission between invasive and native species. Importantly, no *CyHV-3* was detected. This study shows that freshwater fish harbour a high diversity and abundance of viruses, that viruses have likely been associated with fish for millennia, and that there is likely little direct virus transmission between introduced and native species.

## Introduction

Anthropogenic stressors such as pollution, climate change and the introduction of exotic species continue to pose a significant threat to freshwater habitats, with almost one third of all fish species threatened by extinction [1]. The Murray-Darling Basin, the largest freshwater river system in Australia, harbours at least 12 exotic freshwater fish species [2]. Key among these are eastern mosquitofish (*Gambusia holbrooki*), redfin perch (*Perca fluviatilis*) and, most notably, common carp (*Cyprinus carpio*) [2]. Common carp (also known as European carp) were initially introduced into Australia during the mid-1800s for aquaculture operations and again on several occasions throughout the 1900s [3, 4]. During extensive flooding events during the 1970s, carp spread across much of the basin and now represent up to 90% of total fish biomass in the basin [4].

The invasion of carp has been hugely detrimental to Australian freshwater ecosystems [4]. Impacts include increased water turbidity, decreased light penetration, erosion of riverbanks, changes in the abundance and diversity of native invertebrate communities [3, 4, 5] and outcompeting native fish species for habitat and resources [2]. Several control methods have been proposed to control invasive carp; nevertheless, their resilience and high fecundity create significant challenges [6]. This has stimulated research into biological control methods, such as deployment of the virus *cyprinid herpesvirus 3 (CyHV-3)* [7, 8].

*CyHV-3* is a double-stranded DNA virus (family *Alloherpesviridae,* order *Herpesvirales)* first isolated from farmed carp in the late 1990s [9]. Since its discovery, it has been responsible for large disease outbreaks worldwide with a mortality rate of up to 80% in domestic carp [10]. CyHV-3 is transmitted horizontally through direct contact with skin lesions or secretion of viral particles in freshwater where it can survive for up to three days [11]. The host range of CyHV-3 is currently limited to koi and common carp [9]. While *CyHV-3* DNA has been identified in goldfish *(Carrasius auratus)* [12], it is still relatively unclear whether infection occurs in these species [14, 15].

Initial laboratory trials suggest that *CyHV-3* is safe for non-target species [7]. However, little is known about the viruses that naturally circulate in Australian native freshwater fauna, including any prior evidence for the existence of *CyHV-3* [13], nor on the time-scales and frequency with which viruses jump between fish hosts. To completely assess the safety and efficacy of any virus biocontrol agent, including *CyHV-3,* a comprehensive assessment of the viruses that naturally infect both native and invasive species is required.

Following the advent of meta-transcriptomic sequencing, it is now possible to characterise the entire set of viruses – the virome – within a given host [16, 17]. Fish, in particular, harbour a high abundance and diversity of viruses often with deep evolutionary histories [18, 19]. However, despite the antiquity and diversity of fish viruses, there are few studies of virus diversity and evolution in wild freshwater fish populations, particularly in the context of biological invasions.

Determining the viromes of invasive freshwater fish like the common carp will enhance our understanding of the broad-scale factors that influence virus emergence and evolution. As the date and site of their introduction is well-documented in Australian waters, these species can potentially provide important information on the both rate of cross-species transmission and how frequently viruses might move between invasive and native species. In addition, despite representing a small fraction of the earth’s surface water, freshwater environments serve as a habitat for 40-50% of total fish species, harbouring the greatest biodiversity per land area [20]. Such habitats are subject to rapid environmental change, which may significantly impact species connectivity [21]. Since contact and exposure between hosts are vital for cross-species transmission of viruses [22], these species may also inform us on the ecological factors that impact virome composition within a given host.

Herein, we performed a meta-transcriptomic viral survey of invasive and native freshwater fish species across the Murray-Darling Basin in Australia to document the diversity and evolution of freshwater fish viromes and, from this, better understand the ecological drivers of virus evolution and emergence. To the best of our knowledge this is the largest survey of freshwater fish viruses undertaken to date. In particular, we aimed to determine whether *CyHV-3* is already present in common carp in Australia [13], and whether there is evidence for transmission of existing viruses between exotic and native species. As such, we provide important information on the ecological and evolutionary context for the potential release of future virus biocontrols.

## Methods

### Ethics

Fish sampling was conducted with animal ethics approval (ref: 2019/035) from the Animal Ethics Committee (AEC) at Macquarie University, Sydney, NSW. Biosafety was approved by Macquarie University (ref: 5201700856).

### Sample collection

We compared the viromes of native and invasive fish species occupying different areas across the Murray-Darling Basin, Australia (Figure 1). Sampling occurred between January and March 2020. A total of seven native fish species were collected: bony herring *(Nematalosa erebi),* spangled perch *(Leiopotherapon unicolor),* Australian smelt *(Retropinna semoni),* Murray-Darling rainbowfish *(Melanotaenia fluviatilis),* flat-headed gudgeon (*Philypnodon grandiceps*), western carp-gudgeon (*Hypseleotris spp.*) and unspecked hardyhead (*Craterocephalus fulvus*). Three species of invasive fish were also collected: common carp (*Cyprinus carpio*), goldfish (*Carassius auratus*) and eastern mosquitofish *(Gambusia holbrooki).* All fish caught were apparently healthy, with no signs of disease. Fish were caught using boat electrofishing, euthanized and dissected immediately upon capture. Tissue specimens (liver and gills) were placed in RNALater and stored in a portable −80°C freezer, then later in a −80°C freezer in the laboratory at Macquarie University, Sydney. Tissue selection was based on previous studies [18, 37, 44], which show that liver and gill tissue serve as a rich source of viruses. To facilitate virus discovery, multiple individuals (1 – 10) were pooled according to species and the location in which they were captured (SI Table 2; Figure 1).

**Figure 1.**
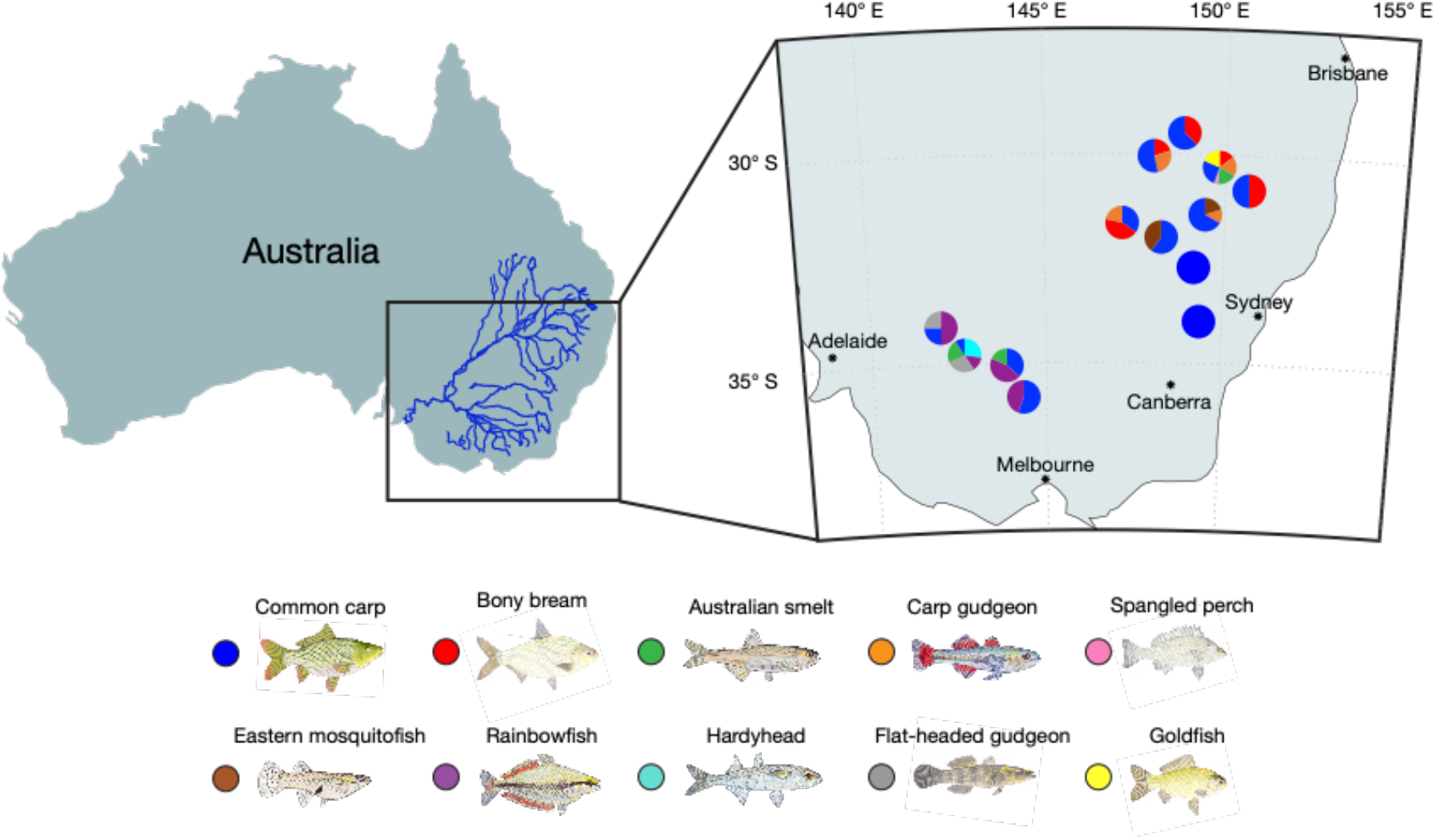
Fish sampling locations. Map illustrating freshwater sampling locations across the Murray-Darling Basin, Australia. Pie-charts show the abundance and diversity of fish species captured at each site. Colours correspond to the fish species sampled as illustrated below the map.

### Total RNA extraction and transcriptome sequencing

Frozen samples of liver and gill tissue were placed together in 600μl of lysis buffer containing 0.5% foaming reagent (Reagent DX, Qiagen) and 1% of ß-mercaptoethanol (Sigma-Aldrich). Submerged tissue samples were homogenized with TissueRuptor (Qiagen) for one minute at 5000 rpm. To further homogenise tissue samples and remove tissue residues, the homogenate was centrifuged at full speed for three minutes. The homogenate was carefully removed and RNA from the clear supernatant was extracted using the RNeasy Plus Mini Kit (Qiagen, Hilden, Germany) following the manufacturer’s protocol. Extracted RNA was quantified using NanoDrop (ThermoFisher) and RNA from each species was pooled corresponding to the site in which they were captured, resulting in 36 sample libraries (SI Table 2). RNA libraries were constructed using the Truseq Total RNA Library Preparation Protocol (Illumina). To enhance viral discovery and reduce the presence of non-viral reads, host ribosomal RNA (rRNA) was depleted using the Ribo-Zero-Gold Kit (Illumina) and paired-end sequencing (150 bp) was performed on the NovaSeq 500 platform (Illlumina). Sample library construction, rRNA depletion and RNA sequencing were performed at the Australian Genome Research Facility (AGRF).

### Virus discovery

Raw Illumina sequence reads (forward and reverse) were initially quality trimmed with Trimmomatic v.0.39 [23] then assembled into contigs *de novo* using Trinity RNA-seq v.2.8.5 [24], with the default parameter settings. Assembled contigs were annotated and compared against the NCBI nucleotide (nt) and non-redundant protein (nr) databases with an e-value threshold of 1×10^-5^ using BLASTn and Diamond (BLASTX) [25]. To initially distinguish between invertebrate and vertebrate-associated viruses, contigs that matched viral sequences were inspected using Geneious v.11.1.5 [26] and translated into amino acid sequences. Amino acid sequences were then used as a single query in additional sequence comparisons against the NCBI nt and nr databases using BLAST algorithms. This method was also used to remove false positives (e.g. host genes and endogenous viral elements) from our analyses. To help exclude instances of index hopping, viral sequences that were identified in multiple libraries were also inspected using Geneious Prime (www.geneious.com) and amino acid pairwise alignments between viral sequences were performed with Multiple Alignment using Fast Fourier Transform (MAFFT) v.7.450 [27], using the E-INS-i algorithm. Abundances of identical viral transcripts were then calculated (see below) and sequences that were present at frequency of <1% of that of the number of reads present in the dominant library were excluded. To determine whether a virus was novel, we followed the broad criteria specified by The International Committee on Taxonomy of Viruses (ICTV) (http://www.ictvonline.org/).

### Inferring the evolutionary history of novel viral sequences

To determine the evolutionary history of the viruses identified in this study and further distinguish between vertebrate and invertebrate-associated viruses (which are usually phylogenetically distinct), we estimated phylogenetic trees using amino acid sequences of stable genomic regions such as RNA-dependant RNA polymerase (RdRp) or DNA polymerase for DNA viruses. To this end, we combined our sequences with background sequences for each respective virus family taken from NCBI/GenBank. Amino acid sequences were aligned with MAFFT v.7.450 [27] using the E-INS-i algorithm. To remove ambiguous regions in the sequence alignment, amino acid sequences were trimmed using trimAl v.1.2 [33]. To estimate phylogenetic trees, selection of the best-fit model of amino acid substitution was determined using the Akaike information criterion (AIC), corrected Akaike information criterion (AICc), and the Bayesian information criterion (BIC) with the ModelFinder function (-m MFP) in IQ-TREE [34, 35]. Sequence data were analysed using a maximum likelihood (ML) approach in IQ-TREE, with 1000 bootstrap replicates. Phylogenetic trees were annotated with FigTree v.1.4.2. and further edited using Adobe Illustrator (https://www.adobe.com).

### Virome composition

To quantify the relative abundance of viral transcripts within the host transcriptome, the RNA-Seq by Expectation (RSEM) value was estimated using Trinity [24], and raw counts from each transcript were standardised against the total number of reads within the given sequencing library. We also used this approach to estimate the relative abundance of a host reference gene, ribosomal protein S13 (RPS13), that is stably expressed in fish. To assess any differences in virome composition between hosts and sites, we calculated alpha diversity (virome richness and Shannon diversity) using Rhea packages [28]. Generalised linear models (GLM) were used to identify the impact of host taxonomy (i.e. species), host geography (i.e. site), water temperature, water pH, water turbidity and species origin (i.e. invasive or native) on both vertebrate-associated virus composition (abundance, richness and diversity) as well as those viruses likely associated with non-fish hosts: the latter should not be affected by aspects of fish biology and hence effectively constitute a negative control. All GLM models were tested using a likelihood-ratio test (χ2) and a Tukey’s post hoc analysis (glht) was performed using the *multcomp* package [29]. To assess viral diversity between samples, we calculated beta diversity using a Bray Curtis dissimilarity with the *phyloseq* package [30]. Differences in virome composition between native and invasive species were calculated using permanova (Adonis test), with the *vegan* package [31]. All statistical analyses were carried out on RStudio V1.2.1335 and plotted using the *ggplot2* package [32].

## Results

We characterised the viromes of ten freshwater ray-finned fish species across seven taxonomic orders (two invasive and five native) at 13 locations across the Murray-Darling Basin in Australia. Total RNA-sequencing was performed on 36 libraries, resulting in a median of 76,528,534 (range 66,015,138 – 95,168,951) reads per library. *De novo* assembly of the sequencing reads resulted in a median of 617,588 contigs (range 198,446 – 1,989,596) per library, with a total of 23,976,218 contigs generated. Analysis of the host reference gene, RPS13, revealed abundances of 0.000001 – 0.0002%, suggesting an inconsistent sequence coverage across all RNA libraries (Figure 2).

**Figure 2.**
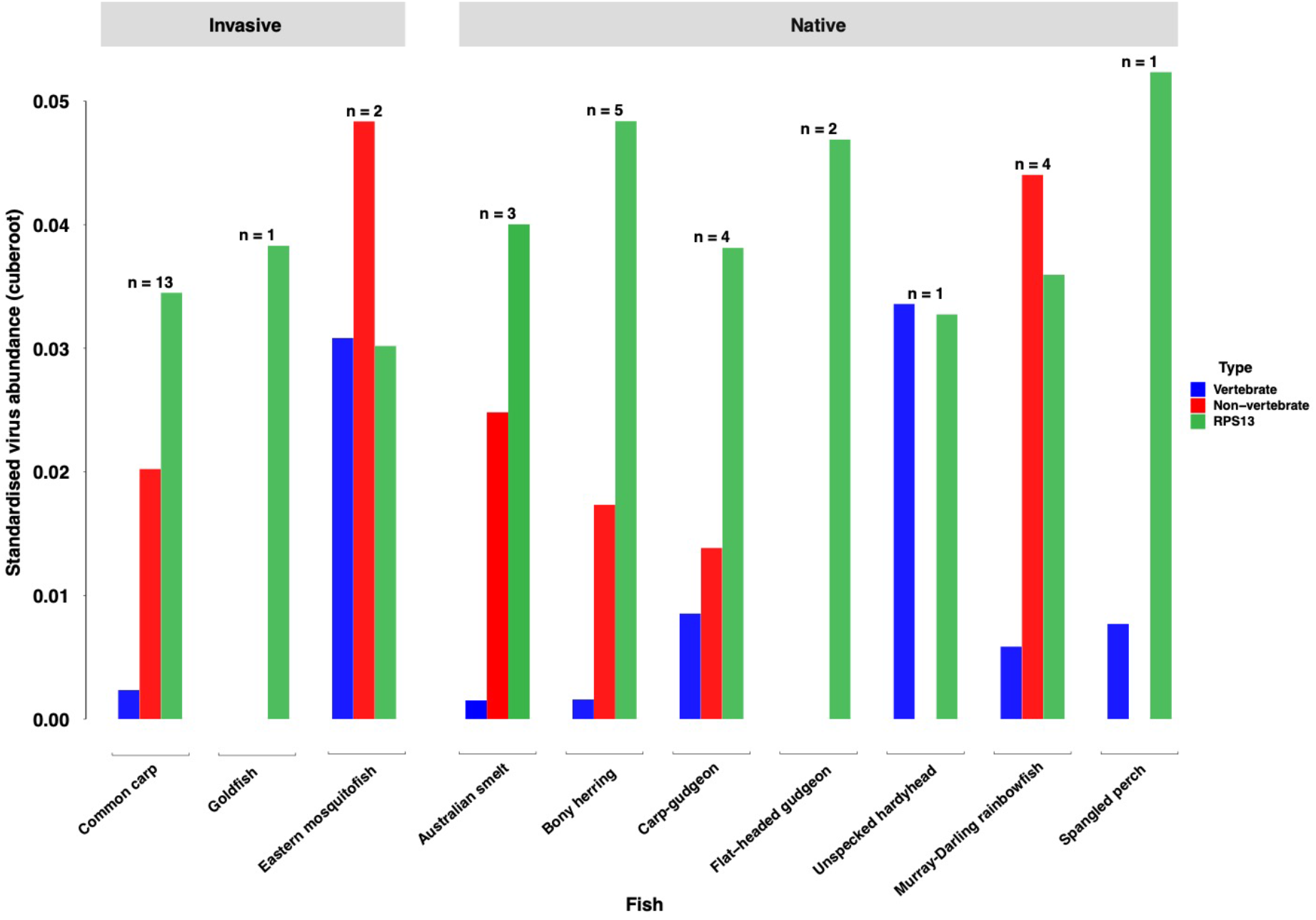
Mean standardised viral abundance across all libraries. Clustered bar chart reveals differences in viral abundance between invasive and native fish species. Blue bars represent vertebrate-associated viruses; red bars represent non-vertebrate associated viruses; and green bars represent host reference gene RPS13. Number of sequencing libraries for each fish species is displayed above bars.

### Abundance and diversity of viruses

We identified 18 viral sequences that were associated with vertebrate hosts and a further 27 that were likely associated with algae, invertebrates and protists in the freshwater environment (SI Figures 1 and 2). Because such non-vertebrate viruses were likely derived from diet or contamination of gill tissue, we primarily focused on vertebrate-associated viruses.

Among the likely vertebrate-associated viruses, we identified viral sequences from 14 viral families. With the exception of a novel poxvirus (family *Poxviridae),* a double-stranded DNA virus, all the viruses identified possessed RNA genomes. The most abundant vertebrate-associated viral transcripts were those assigned to the *Arenaviridae* (49% of all vertebrate-associated viruses), *Hepeviridae* (20%), *Chuviridae* (21 %), *Astroviridae* (3%) and *Flaviviridae* (2%) families. Other likely vertebrate viral transcripts detected were assigned to the *Coronaviridae* (<1 %) *Caliciviridae* (<1 %), *Picornaviridae* (<1 %), *Paramyxoviridae* (<1 %), *Hantaviridae* (<1 %), *Bornaviridae* (<1 %), *Poxviridae* (<1 %), *Reoviridae* (<1 %) and *Rhabdoviridae* (<1 %) families. The most common vertebrate-associated viruses identified were astroviruses, detected in three host species (eastern mosquitofish, Murray-Darling rainbowfish, spangled perch). In addition, arenaviruses were detected in two host species (western carp-gudgeon, eastern mosquitofish) along with hepeviruses (common carp, eastern mosquitofish). All other viruses were identified in one host species (Figure 3).

**Figure 3.**
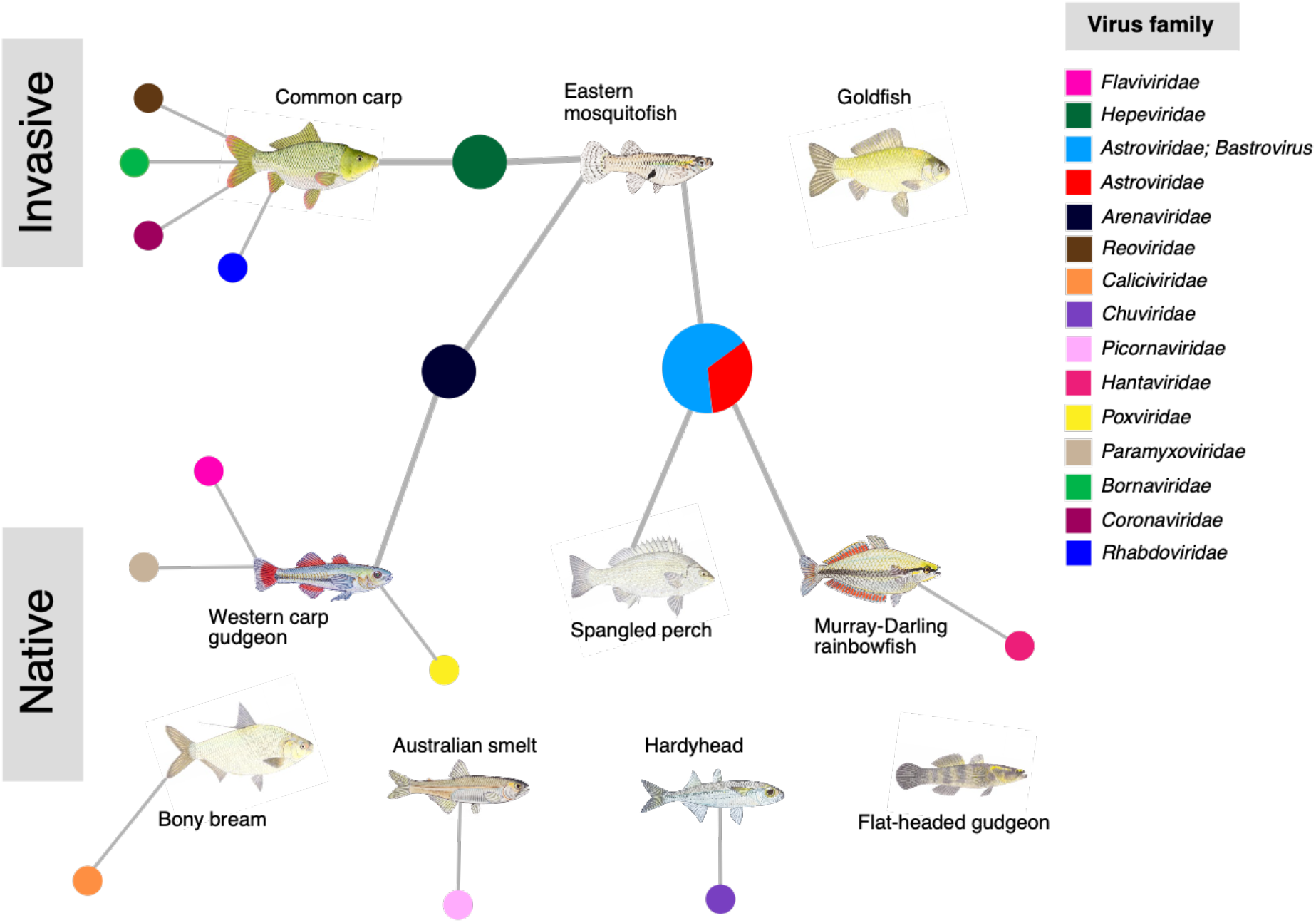
Network diagram displaying vertebrate-associated viruses identified in native and invasive freshwater fish. Colours of each node represents a virus family. Both goldfish and flat-headed gudgeon contained no vertebrate-associated viruses.

Among the viruses likely associated with non-vertebrate hosts (i.e. those infecting arthropods, fungi, plants and protozoans), a large proportion (70%) were unclassified, comprising picorna-like viruses, rhabdo-like viruses, tombus-like viruses and narna-like viruses [45] (SI Figure 1). We also detected viral transcripts that could be assigned to the *Nodaviridae* (27.1 %), *Permutotetraviridae* (1.3%), *Dicistroviridae* (1.2%) and *Phenuiviridae* (<1 %) families. Although viruses within the *Nodaviridae* have been shown to infect fish [46], all of the nodavirus sequences identified here clustered with viruses from invertebrate hosts (SI Figure 2), strongly suggesting they were similarly associated with fish diet or contamination of gill tissue.

### Phylogenetic relationships of vertebrate-associated viruses

To infer the phylogenetic relationships and hence the evolutionary history of the viruses newly identified here, we focused on stable genomic regions such as the RdRp in RNA viruses and DNA polymerase in the case of the novel poxvirus. Using these genomic regions, we identified seven negative-sense single-stranded RNA (-ssRNA) viruses (families *Arenaviridae, Bornaviridae, Chuviridae, Hantaviridae, Paramyxoviridae, Rhabdoviridae*), nine positive-sense single-stranded RNA (+ssRNA) viruses (families *Astroviridae, Caliciviridae, Coronaviridae, Flaviviridae, Hepeviridae, Picornaviridae),* one double-stranded RNA (dsRNA) virus (family *Reoviridae)* and one double-stranded DNA (dsDNA) virus (family *Poxviridae).* We now describe each of these groups in turn.

### Negative-sense single-stranded (-ssRNA) viruses

We identified -ssRNA viruses that occupied phylogenetic positions that were broadly indicative of long-term virus-host co-divergence, with many fish viruses falling basal to reptile and mammalian viruses (Figure 4). Notably, we identified two novel arenaviruses that clustered with members of the newly formed *Antennavirus* genus that includes fish hosts [18, 36]. *Western carp-gudgeon arenavirus,* found at Narrabri creek shared 36.9% amino acid sequence similarity with its closest relative, *Wenling frogfish arenavirus 1* [18] and grouped with recently discovered arenaviruses from pygmy gobies (*Eviota zebrina*) sampled from an Australian coral reef [37]. *Eastern mosquitofish arenavirus* found in the Macquarie River shared 84.5% amino acid sequence similarity with its closest available relative, *Wenling frogfish arenavirus 1* [18].

**Figure 4.**
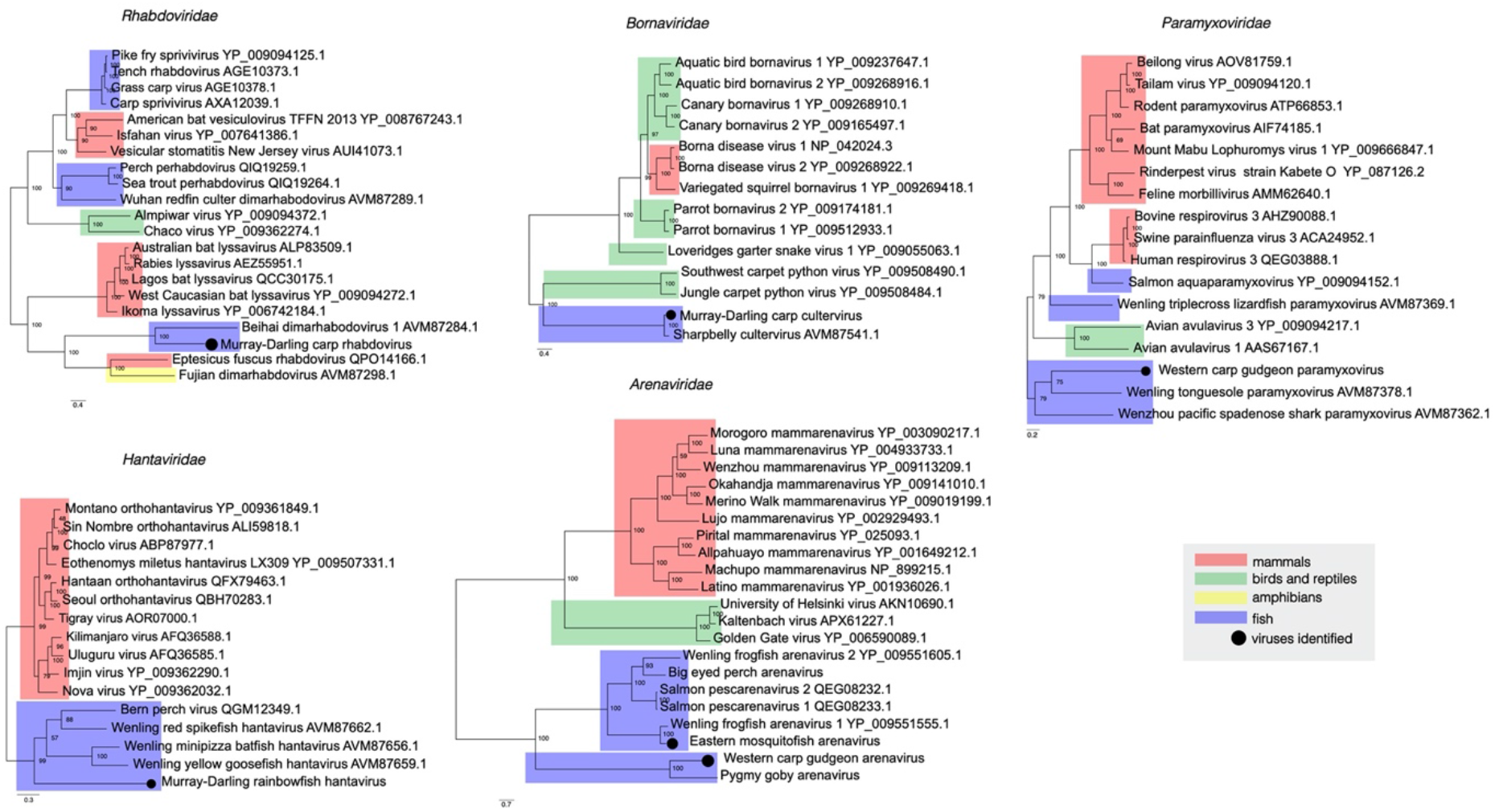
Phylogenetic relationships of negative-sense single-stranded vertebrate-associated viruses identified in this study. Viruses identified here are shown as a black circle. Maximum likelihood trees were estimated using amino acid sequences of the RdRp gene and mid-point rooted for clarity only. Bootstrap values are represented as a percentage with branches scaled to amino acid substitutions per site. Tree branches are highlighted to represent host class: red, mammals; green, birds and reptiles; yellow, amphibians; and blue, fish. Tip labels represent virus name and NCBI/GenBank accession numbers.

The divergent hantavirus detected in the Murray-Darling rainbowfish falls basal to mammalian hantaviruses (orthohantaviruses) and clustered with members of the *Actantavirinae* and *Agantavirinae* subfamilies that include ray-finned and jawless fish hosts [18, 36] (Figure 4). This virus had only 27.3% amino acid similarity with its closest relative, *Bern perch virus* (NCBI/GenBank: QGM12349.1). Broad patterns of virus-host codivergence can similarly be seen in the cultervirus identified in carp from Lake Burrendong. BLAST analysis identified *sharpbelly cultervirus* [18] as the closest relative of all genomic regions, including the L gene (93% amino acid similarity), glycoprotein (86.3%) and nucleoprotein (92.9%).

Our virological survey revealed the complete genome of a novel chuvirus in the unspecked hardyhead in the Edward River. This *Hardyhead chuvirus* displayed three open reading frames (ORFs), representing the L protein (RdRp), glycoprotein and nucleoprotein. Our analysis identified *Guangdong red-banded snake chuvirus* [18] as the closest relative of the L protein (44% amino acid similarity), *Wenling fish chu-like virus* [18] as the closest relative of the glycoprotein (41%), and *Herr Frank virus 1* [41] as the closest relative of the nucleoprotein (34%). *Hardyhead chuvirus* formed a distinct phylogenetic clade with all other vertebrate-associated chuviruses (Figure 5).

**Figure 5.**
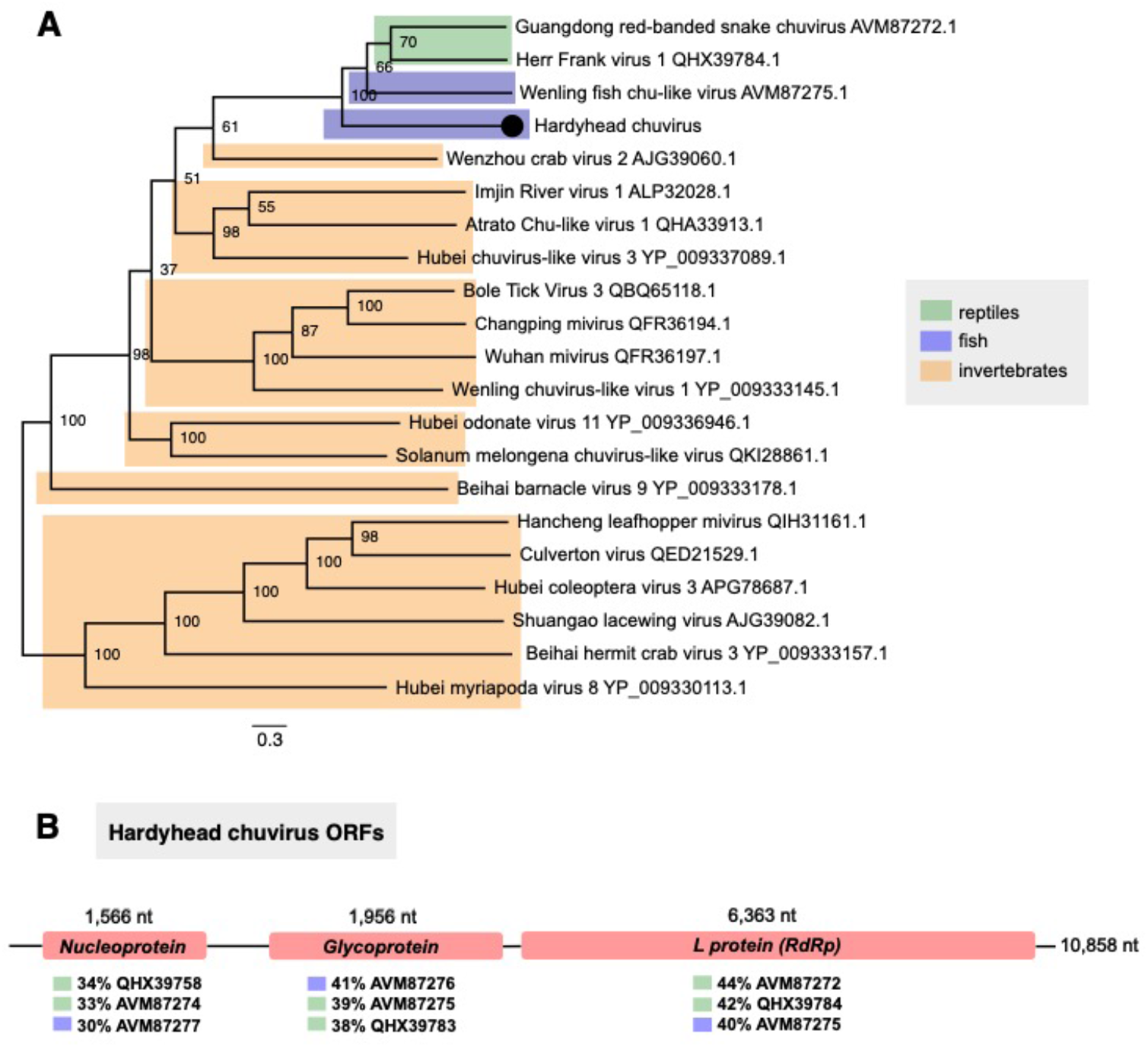
Phylogenetic relationships and genome organisation of *hardyhead chuvirus.* (A) Phylogenetic relationships between viruses within the *Chuviridae.* Novel chuvirus identified in the unspecked hardyhead is represented as a black circle. The phylogenetic tree was estimated using amino acid sequences of the RdRp gene and midpoint rooted for clarity only. The scale bar represents amino acid substitutions per site. Bootstrap values are shown as a percentage. Tree branches are highlighted to distinguish between vertebrate and invertebrate-associated viruses: green, reptiles; blue, fish; and orange, invertebrates. Tip labels represent virus name and NCBI/GenBank accession numbers. (B) Genome organisation of *hardyhead chuvirus*. Diagram illustrates the structure and length of each genomic segment. Percentages below each segment reveal the closest relatives (NCBI/GenBank accession number), with the L protein more related to reptile chuviruses; glycoprotein more related to fish chuviruses; and nucleoprotein more related to reptile chuviruses.

We also detected a novel paramyxovirus in western carp-gudgeon in the Bogan River. This divergent viral sequence shared 35.2% L gene amino acid similarity with its closest relative, *Wenling tonguesole paramyxovirus* (genus *Cynoglossusvirus*, family *Paramyxoviridae*) [18]. These viruses grouped with *Wenzhou pacific spadenose shark paramyxovirus* (genus *Scoliodonvirus),* together falling basal to other members of the *Paramyxoviridae* family. In addition, a novel rhabdovirus in common carp similarly formed a distinct clade, basal to other fish-infecting rhabdoviruses. This virus shared 35.7% amino acid L gene sequence similarity with *Beihai dimarhabdovirus* that was also identified in fish [18] and clustered with other dimarhabdoviruses, including those found in the spotted paddle-tail newt from China [18] and the big brown bat *(Eptesicus fuscus)* from the USA (NCBI/Genbank: QPO14166.1). Across all genera within the *Rhabdoviridae,* lyssaviruses were the closest relatives to this clade (Figure 4), with *Murray-Darling carp rhabdovirus* sharing 31.6% amino acid L gene similarity with *rabies lyssavirus* [43].

### Positive-sense single-stranded (+ssRNA) viruses

We identified a viral sequence in common carp that shared 50.7% RdRp sequence similarity with *Pacific salmon nidovirus* (family *Coronaviridae)* [60]. *Murray-Darling carp letovirus* also exhibited sequence similarity (46.2%) with gammacoronaviruses, including *bottlenose dolphin coronavirus* and *beluga whale coronavirus* [58, 59]. This virus grouped with both *Pacific salmon nidovirus* and *Microhyla letovirus* [81], which together form an outgroup to all other coronaviruses (Figure 6).

**Figure 6.**
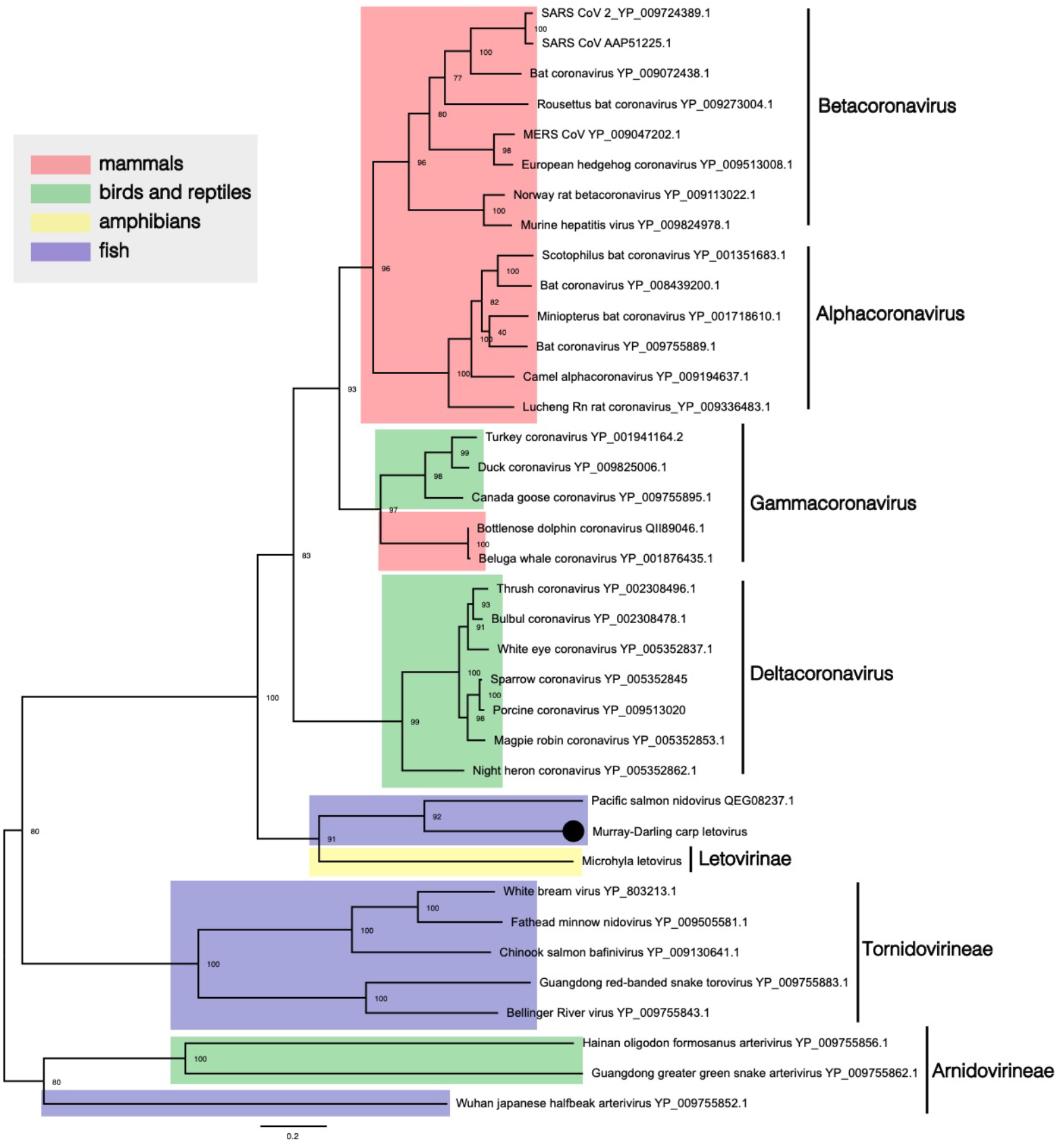
Phylogenetic relationships within the virus order *Nidovirales.* The novel letovirus detected in common carp is labelled with a black circle. The phylogenetic tree was estimated using the orf1ab polypeptide (polymerase associated) and midpoint rooted for clarity alone. Scale bar represents amino acid substitutions per site. Bootstrap values are shown as a percentage. Taxon labels are highlighted: blue, fish; red, mammals; green, birds and reptiles; yellow, amphibians. Virus taxonomic names are labelled to the right. Tip labels represent virus name and NCBI/GenBank IDs.

We also identified a novel flavivirus (genus *Flavivirus,* family *Flaviviridae)* in western carp-gudgeon in the Bogan River. This viral sequence exhibited 33-36% NS5 amino acid sequence similarity with its closest relatives, *Cyclopterus lumpus virus* [38], *Tamana bat virus* [39], *salmon flavivirus* [40] and *Wenzhou shark flavivirus* [18]. All these viruses fall basal to vector-borne viruses within the genus *Flavivirus* (Figure 7).

**Figure 7.**
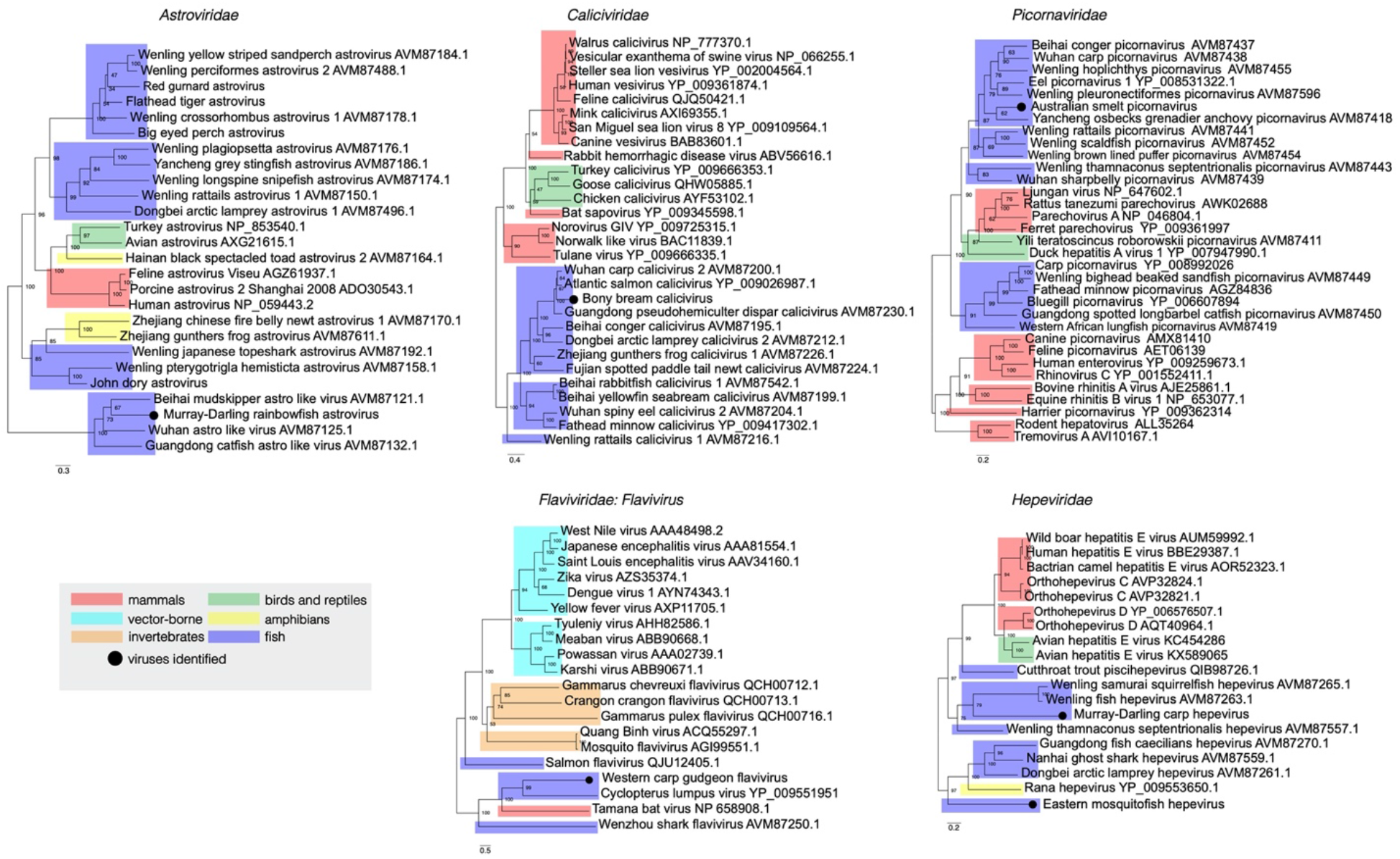
Phylogenetic relationships of positive-sense singe-stranded vertebrate-associated viruses identified in this study. Viruses identified here are shown as a black circle. Maximum likelihood trees were estimated using amino acid sequences of the RdRp gene and NS5 gene for the novel flavivirus. Trees were mid-point rooted for clarity only and bootstrap values are represented as a percentage with branches scaled to amino acid substitutions per site. Tree branches are highlighted to represent host class: red, mammals; cyan, vector-borne viruses; green, birds and reptiles; yellow, amphibians; blue, fish; and orange, invertebrates. Tip labels represent virus name and NCBI/GenBank accession IDs.

Among other positive-sense RNA viruses identified, a novel astrovirus, calicivirus and picornavirus all grouped with other fish hosts and expanded the phylogenetic diversity of these virus families (Figure 7). The novel astrovirus identified in Murray-Darling rainbowfish shared 40% RdRp amino acid similarity with *Wuhan astro-like virus* [18]. This virus clustered with other astro-like viruses discovered in fish, including *Beihai mudskipper astro-like virus* [18] and *Guangdong catfish astro-like virus* [18]. This was similarly observed in *Australian smelt picornavirus*, which clustered with picornaviruses found in other freshwater fish, including those from eels (*Anguilla anguilla*) [48] and carp sampled from China [18]. The *Caliciviridae* includes two genera that infect fishes: saloviruses associated with salmonid hosts and minoviruses associated with cyprinid hosts [49]. Recently, several caliciviruses have been discovered in ray-finned and jawless fish [18, 50]. The novel calicivirus identified in bony herring expands the diversity of fish viruses as it shared 80% amino acid similarity with *Atlantic salmon calicivirus* [50] and clustered with other freshwater fish caliciviruses, including *Wuhan carp calicivirus* [18] and *Guangdong pseudohemiculter dispar calicivirus* [18].

### Double-stranded RNA (dsRNA) viruses

We identified a novel dsRNA virus in carp in the Castlereagh River that could be assigned to the *Reoviridae.* This divergent virus shared 40% RdRp amino acid similarity with its closest relative, *Wenling scaldfish reovirus* [18], together forming a clade basal to the genus *Aquareovirus* that are known to cause considerable disease in some fish species [42] (Figure 8).

**Figure 8.**
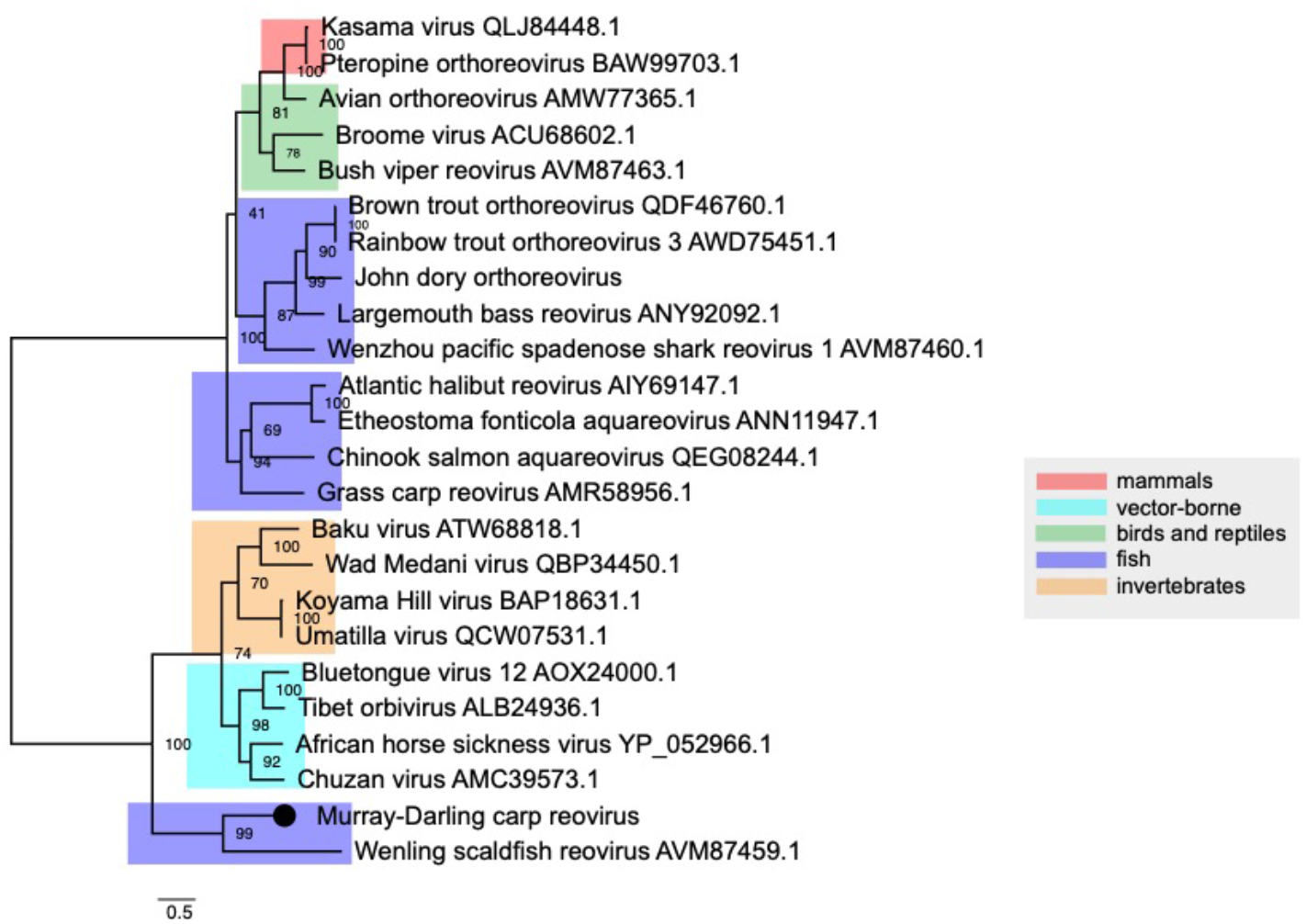
Phylogenetic relationships within the *Reoviridae.* Novel reovirus identified here shown as a black circle. Maximum likelihood tree was estimated using amino acid sequences of the RdRp gene. Tree was mid-point rooted for clarity only and bootstrap values are represented as a percentage with branches scaled to amino acid substitutions per site. Tree branches are highlighted to represent host class: red, mammals; cyan, vector-borne viruses; green, birds and reptiles; blue, fish; and orange, invertebrates. Tip labels represent virus name and NCBI/GenBank accession IDs.

### Double-stranded DNA (dsDNA) viruses

A key observation of our study was the absence of Cyprinid herpesviruses, including *CyHV-3,* as well as other *Alloherpesviridae,* in any of the 36 RNA libraries. Similarly, although members of the *Hepadnaviridae* are commonly detected in fish [37, 44, 77, 78] they were notably absent in our samples. The only DNA virus detected in this study was a novel poxvirus *(Poxviridae)* identified in western carp-gudgeon. This virus shared DNA polymerase amino acid sequence similarity with *salmon gill poxvirus (SGPV)* (61 %) [51]. We also detected other genomic regions such as DNA-dependant RNA polymerase subunit rpo22 (49.5%), DNA-dependant RNA polymerase subunit rpo19 (40%), DNA binding virion core protein I1L (28.1%), A16L (32.9%) and SGPV079 (40%) (SI Table 1). Both *SGPV* and *western carp-gudgeon poxvirus* form a highly divergent clade within the subfamily *Chordopoxvirinae* that is strongly indicative of virus-host co-divergence (Figure 9).

**Figure 9.**
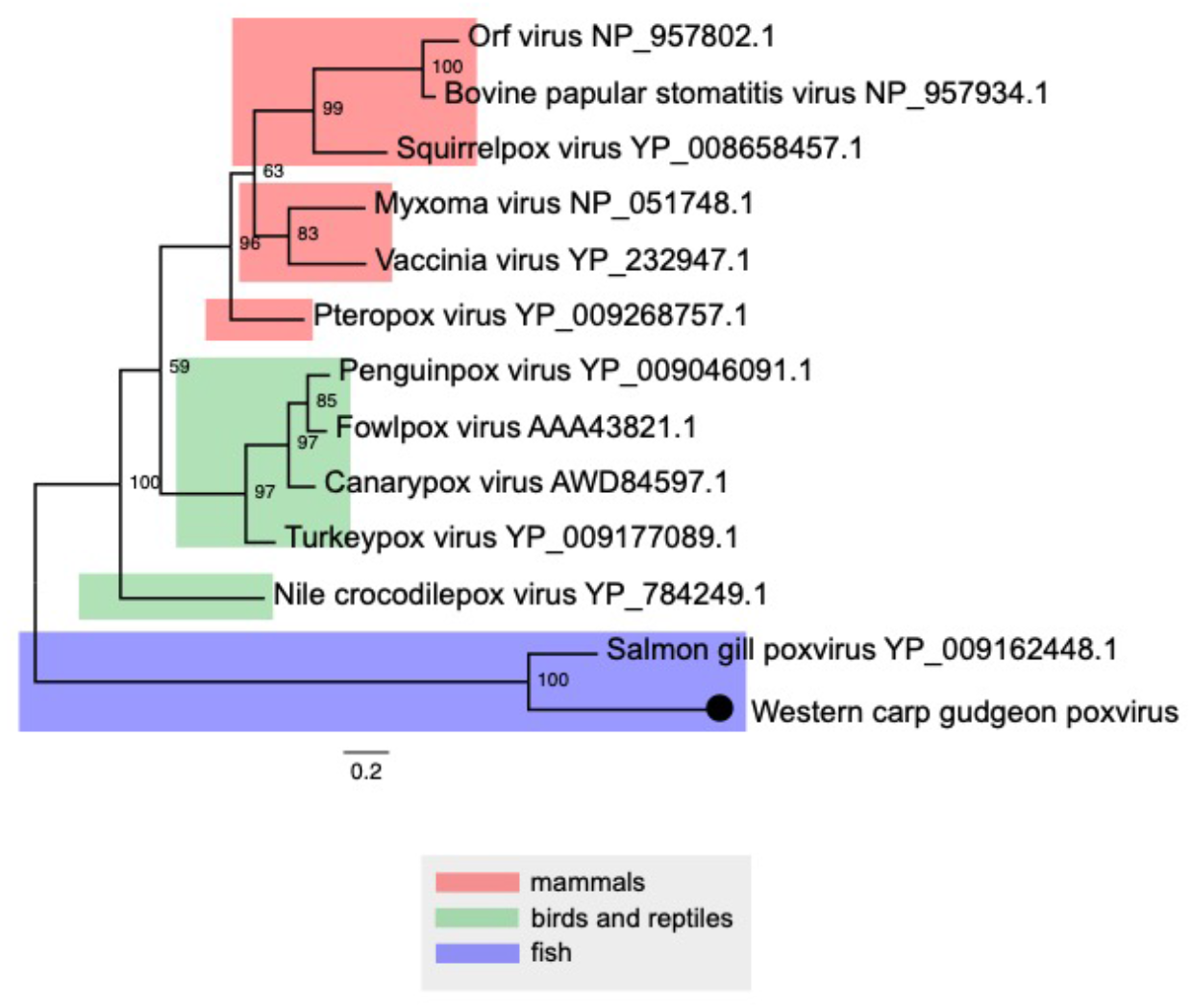
Phylogenetic relationships within the subfamily *Chordopoxvirinae* (family *Poxviridae*). Phylogenetic tree reveals virus-host co-divergence, with fish viruses falling basal to reptilian, avian and mammalian poxviruses. Novel poxvirus identified here shown as a black circle. Maximum likelihood tree was estimated using amino acid sequences of the DNA polymerase gene. Tree was mid-point rooted for clarity only and bootstrap values are represented as a percentage with branches scaled to amino acid substitutions per site. Tree branches are highlighted to represent host class: red, mammals; green, birds and reptiles; blue, fish. Tip labels represent virus name and NCBI/GenBank accession IDs.

### Virome composition, ecological and environmental factors

We next examined whether and how vertebrate virome composition in a range of native and introduced Murray-Darling Basin fish was associated host ecological factors, namely host species, geography (i.e. sampling site), water quality (temperature, pH and turbidity). GLMs revealed host species (χ2= 7.5^-6^, df= 9, p = 0.001) as the best predictor of viral abundance (i.e. the standardised number of viral sequencing reads) (Figure 2). In particular, the eastern mosquitofish had significantly higher viral abundance compared to Australian smelt (Tukey: z=3.976, p=0.002), bony herring (Tukey: z=4.334, p=0.001), western carp-gudgeon (Tukey: z=4.019, p=0.002), common carp (Tukey: z=4.665, p=0.001), flat-headed gudgeon (Tukey: z=3.632, p=0.001), goldfish (Tukey: z=3.632, p=0.001) and rainbowfish (Tukey: z=4.110, p=0.001). However, the high viral abundance in the eastern mosquitofish was driven by one sample containing an extremely high abundance of arenaviruses, accounting for 76% of its total vertebrate virome and 49% of all vertebrate-associated viral reads. We found no evidence for an association between viral abundance and host geography (p = 0.111), water turbidity (p = 0.804), water temperature (p = 0.709) nor water pH (p = 0.141).

We calculated alpha diversity to assess any differences in virome composition (abundance and diversity) between hosts and sites. This included the observed virus species richness (the number of viruses found in each sequencing library) and Shannon diversity (both the number of viral families and abundance of viral reads in a given host). We found no association between observed viral species richness and host species (p = 0.286), host geography (p = 0.748), water turbidity (p = 0.826), water temperature (p = 0.625) and water pH (p = 0.115). Similarly, there was also no observed association between Shannon diversity and host taxonomy (p = 0.117), host geography (p = 0.55), water turbidity (p = 0.546), water temperature (p = 0.206) and water pH (p = 0.039).

### Virome composition of native versus invasive fish species

While carp and native fish species were sampled together at 10 out of 13 sites (Figure 1), they shared no vertebrate-associated viruses. Although we identified hepeviruses in common carp and eastern mosquitofish, these were highly divergent and exhibited only 20% amino acid similarity such that they reflect ancient divergence events. These viruses were also distinct in that *Murray-Darling carp hepevirus* clustered with ray-finned fish hosts [18], while *eastern mosquitofish hepevirus* formed a distinct basal clade with amphibian [54], jawless fish and cartilaginous fish hosts [18] (Figure 7).

While the native and invasive fish species largely had distinct viromes, two vertebrate-associated virus families were present in both: arenaviruses (western carp-gudgeon and eastern mosquitofish) and bastroviruses (spangled perch and eastern mosquitofish) (Figure 3). However, the arenaviruses identified in the eastern mosquitofish and western carp-gudgeon were highly divergent, exhibiting only 27.9% amino acid similarity, with *western carp-gudgeon arenavirus* falling basal to *eastern mosquitofish arenavirus* (Figure 4). The bastroviruses detected in eastern mosquitofish and spangled perch shared 57.1% RdRp amino acid similarity and formed a distinct clade with other bastrovirus sequences identified in *Culex* mosquitos [52], bats (NCBI/GenBank: NC_035471.1) and sewage samples in Brazil [53] (SI Figure 3). Because bastroviruses have genomic features that resemble hepeviruses [54], both these contigs had matches to invertebrate and vertebrate-associated hepeviruses such that their true hosts could not be easily determined.

We also examined whether there were any differences in alpha and beta diversity between native and invasive freshwater fishes. Accordingly, we found no association between host origin (i.e. invasive or native) and virome abundance (p = 0.390). When assessing alpha diversity, we similarly found no association between host origin and virome richness (p = 0.626) nor Shannon diversity (p = 0.425). Likewise, this result was also observed when examining beta diversity (p = 0.602).

### Associations between host ecology and non-vertebrate viruses

To assess any associations between host ecology and non-vertebrate viruses, we similarly performed GLMs using the aforementioned ecological factors as a negative control. As expected, this revealed no association between non-vertebrate viral abundance and host taxonomy (p = 0.200), host geography (p = 0.101), host origin (p = 0.998), water turbidity (p = 0.421), water temperature (0.282) and water pH (p = 0.343). We similarly found no association between non-vertebrate virome richness and host taxonomy (p = 0.204), host geography (p = 0.090), host origin (p = 0.675), water turbidity (p = 0.398), water temperature (p = 0.072) and water pH (p = 0.461). We also found no evidence for an association between Shannon diversity and host taxonomy (p = 0.691), host geography (p = 0.173), host origin (p = 0.876), water turbidity (p = 0.571), water temperature (p = 0.334) and water pH (p = 0.578). This was also observed when assessing statistical associations between beta diversity and host species (p = 0.684), host origin (p = 0.239) and host geography (p = 0.501).

## Discussion

Our meta-transcriptomic viral survey of native and invasive fish across the Murray-Darling Basin, Australia, revealed a high diversity and abundance of viruses, including the identification of 45 novel virus species that infected seemingly healthy fish or nonvertebrate hosts in the freshwater environment. Crucially, however, we observed no clear examples of recent cross-species transmission among fish hosts, particularly between invasive and native species, nor any evidence for the presence of *CyHV-3* from a total of 36 RNA sequencing libraries. Hence, these data provide further evidence of the absence of *CyHV-3* in Australia [7]. Similarly, our analysis failed to detect other cypriniviruses (i.e. *CyHV-1*, *CyHV-2),* despite previous reports of the presence of *CyHV-2* in the Murray-Darling Basin [75, 76].

The only instance of co-occurrence of viruses from the same family in both invasive and native species were the presence of arenaviruses in native carp-gudgeon and invasive mosquitofish. However, these viruses were so divergent that they likely represent ancient common ancestry rather than recent cross-species transmission (Figure 4). Eastern mosquitofish, introduced into Australia during the early 1920s to control mosquito populations [61], are now widespread across the Murray-Darling Basin and have become a successful invasive species [62]. Their abundance primarily impacts smaller native fish (such as carp-gudgeon, rainbowfish and hardyheads) since they typically outcompete these species and disrupt food webs [62]. As well as being highly divergent, *western carp-gudgeon arenavirus* formed a basal clade with a recently discovered arenavirus in the pygmy goby sampled from an Australian coral reef [37], a member of the same fish order (*Gobiiformes*). These data further suggest that arenaviruses may have been circulating in Gobiiform fishes (gobies and gudgeons) in Australia prior to the introduction of eastern mosquitofish.

We also identified a novel coronavirus - *Murray-Darling carp letovirus* - that shared 50.7% amino acid sequence similarity with *Pacific salmon nidovirus* and 46.2% amino acid similarity with gammacoronaviruses. The *Coronaviridae* (order *Nidovirales)* can be split into two subfamilies: the *Orthocoronavirinae,* associated with birds and mammals and the *Letovirinae* associated with amphibians [57]. That both *Murray-Darling carp letovirus* and *Pacific salmon nidovirus* formed a sister clade to only member of the *Letovirinae* subfamily, *Microhyla letovirus* (genus *Alphaletovirus)* identified in the ornamental pygmy frog (*Microphyla fissipes*) (Figure 6) [57], suggests that fish may be common and ancient hosts for the *Letovirinae.* It is also notable that *Murray-Darling carp letovirus* and *Pacific salmon nidovirus* are highly divergent from the other *Nidovirales* that are known to infect fish (e.g. *Chinook salmon bafinivirus*) [56].

The phylogenetic range of the *Chuviridae* largely incorporates invertebrate hosts with diverse genomes (segmented, unsegmented and circular) [64]. Recently, chuviruses have been discovered in vertebrates, all possessing three segments [18, 41]. The novel chuvirus detected here in the unspecked hardyhead displayed these genomic features with the L gene (RdRp), S gene (glycoprotein) and N gene (nucleoprotein) all related to fish and reptile viruses (Figure 5) [18, 41]. The phylogenetic position of this vertebrate clade suggests the ancestors of the viruses may be of invertebrate origin, particularly those that inhabit aquatic ecosystems. For instance, the closest related invertebrate viruses were *Wenzhou crab virus* [64], *Imjin River virus* (mosquitos) [65] and *Atrato chu-like virus* (mosquitos). Similarly, chuvirus endogenous viral elements have been detected in several freshwater fish species [18].

We identified a novel flavivirus in western carp-gudgeon across the Bogan River. This virus falls basal to mammalian vector-borne viruses in phylogenetic trees, grouping with viruses from other vertebrate hosts including *Cyclopterus lumpus virus, Tamana bat virus and Wenzhou shark flavivirus.* Although *western carp-gudgeon flavivirus* was detected in apparently healthy fish, *in vivo* flavivirus replication was recently demonstrated in Chinook salmon (*Oncorhynchus tshawytscha*) that were associated with abnormal mortalities in the Eel River, California [40]. While there is still no clear link between flavivirus infection, transmission and disease in aquatic hosts, these data suggest that flaviviruses may be common in fish species. Moreover, the basal phylogenetic positions of aquatic flaviviruses also suggests that these viruses may be the ancestors of notable vector-borne viruses (Figure 7). Nevertheless, gaps still remain in the evolutionary history of the genus *Flavivirus* and will likely be bridged with additional metagenomic studies.

In broad terms, the evolutionary histories of many vertebrate viral families appear to generally follow patterns of long-term virus-host co-divergence, albeit with regular crossspecies transmission [18,19]. This evolutionary pattern can be observed in the phylogenies of the cultervirus, poxvirus and arenaviruses identified here. The *Bornaviridae* contain three genera with 11 currently classified viral species that infect mammals, birds and reptiles [67]. The only fish virus identified to date falls within the genus *Cultervirus*, comprising *Sharpbelly cultervirus* from China [18]. We identified this virus (i.e. transcripts with 93% L gene amino acid similarly) in common carp in Australia. Intriguingly, both fish hosts are members of the *Cyprinidae* that date as far back as the Cretaceous to Jurassic periods [68, 69]. Recent molecular clock dating using endogenous viral elements also showed that culterviruses likely emerged early on during the course of vertebrate evolution, more than 50 million years ago [18].

Patterns of long-term virus-host co-divergence can also be seen in the evolutionary history of the *Chordopoxvirinae*. *Western carp-gudgeon poxvirus* expands the host range of the *Chordopoxvirinae* subfamily within the *Poxviridae,* forming a highly divergent clade with the only other fish-infecting chordopoxvirus discovered to date, *salmon gill poxvirus (SGPV)* (Figure 9). Since its classification in 2015, several cases of *SGPV* have been identified in farmed salmon with complex gill disease, although the reservoir host is unknown [55]. The phylogeny of the *Chordopoxvirinae* mirrors that of vertebrate hosts, strongly suggesting long-term virus-host co-divergence (Figure 9). Similarly, the phylogeny of the *Arenaviridae* displays a basal fish clade that is characterised by long branches with a large degree of divergence (Figure 4).

On this evolutionary backbone of ancient virus-host co-divergence, we also detected cases of cross-species virus transmission during evolutionary history, although the time-scales of these events are uncertain. For example, we discovered a novel reovirus that formed a basal divergent clade to other fish viruses within the genus *Aquareovirus* (Figure 8). *Murray-Darling carp reovirus* was more closely related to viruses that infect scaldfish [18] rather than other cyprinid hosts, which are highly susceptible to reovirus infection (e.g. 80% mortality in grass carp) [42]. These patterns were also observed in the phylogeny of the *Rhabdoviridae,* with *Murray-Darling carp rhabdovirus* forming a distinct phylogenetic clade with other recently discovered rhabdoviruses in fish, the big brown bat (NCBI/Genbank: QPO14166.1) and the spotted paddle-tail newt [18]. Rhabdoviruses exhibit a very broad host range including invertebrates, plants, mammals, fish, amphibians, birds and reptiles [70]. Notable among the *Rhabdoviridae* are the lyssaviruses that can cause high mortality in human populations (e.g. *rabies lyssavirus).* Intriguingly, *Murray-Darling carp rhabdovirus* and its closest relatives form a sister clade to the genus *Lyssavirus,* suggesting these viruses may have a fish-infecting ancestor (Figure 4).

Although carp are widespread and abundant across the Murray-Darling Basin, they displayed lower viral abundance than some of the other hosts sampled (Figure 2). This could be partly explained by the large phylogenetic distance between carp and other fish in the Murray-Darling Basin. For instance, aside from bony herring, all the fish studied here are members of the *Acanthopterygii* (Percomorpha). This could also explain why invasive mosquitofish harboured similar viruses to native gudgeon species (e.g. arenaviruses).

Although not always the case [22], cross-species virus transmission often occurs between phylogenetically related hosts, particularly those that display conserved cell receptors [66, 71]. In addition, it has been widely suggested that introduced populations are associated with a lower pathogen prevalence and diversity than native species [73, 74]. For example, because invasive species are often established from a small founder population, they likely acquire only a small proportion of pathogens in the novel environment [73, 74]. Once a species rapidly becomes invasive, the diversity of pathogens in this population should remain small, such that the lack of disease likely facilitates the success of invasive species [73, 74].

It is important to note, however, that there were necessary variations within our sampling. For instance, carp and native fish species were sampled together at 10 out of 13 sites, with carp sampled from all 13 sites (Figure 1). In addition, all other fish species were sampled from 1 −5 sites. While an artifact of the distribution of the fishes, such gaps limit the power of our statistical analyses and perhaps prevent the detection of ecological associations on virome composition within host species, including between invasive and native fish. In addition, due to animal ethics constraints, we were limited to only a subset of native fish species. Nevertheless, the native species examined in this study are generally those present in the highest densities.

Finally, our analysis detected no viruses that are listed as reportable notifiable aquatic diseases in the Murray-Darling Basin [72]. Such notifiable aquatic diseases include epizootic haematopoietic necrosis virus (EHNV – *Iridoviridae)* and Spring viraemia of carp (SVC – *Rhabdoviridae).* EHNV is known to cause high-impact infections in redfin perch and is capable of infecting other freshwater fish in the Murray-Darling Basin, including silver perch *(Bidyanus bidyanus),* Macquarie perch *(Macquaria australasica),* Murray-Darling rainbowfish, freshwater catfish (*Tandanus tandanus*) and invasive mosquitofish [47]. Although thought to be endemic to the Murray-Darling Basin (upper Murrumbidgee River), EHNV was last reported in 2012 [63]. Similarly, we did not detect the emerging dwarf gourami iridovirus *(Iridoviridae)* that causes infectious spleen and kidney necrosis in several species of native Australian fish [79, 80].

In sum, our metagenomic surveillance revealed a marked lack of virus exchange between native and invasive fish species in the Murray-Darling Basin, including those viruses found in invasive common carp. At face value these data suggest that there is minimal virus transmission from common carp to native fish species, although more extensive sampling is needed to fully address this issue. By investigating the viromes of native and invasive fishes, we provide the first data on viruses that naturally circulate in a 2,200 km river system, enhancing our understanding of the evolutionary history of vertebrate viruses.

## Data Availability

All sequence reads generated in this project are available under the NCBI Short Read Archive (SRA) under BioProject PRJNA701716 and all consensus virus genetic sequences have been deposited in GenBank under accessions MW645018-MW645046.

## Acknowledgements

This work was funded by a Recreational Fishing Trust grant (SF024) awarded to JLG, JEW and ECH; an ARC Discovery grant (DP200102351) awarded to ECH and JLG; a Macquarie University grant awarded to JLG; and the Department of Biological Sciences at Macquarie University. ECH is funded by ARC Australian Laureate Fellowship (FL170100022). J.L.G. is funded by a New Zealand Royal Society Rutherford Discovery Fellowship (RDF-20-UOO-007). We thank Justin Stanger, Patrick Martin, Karen Scott, Michael Rodgers, Duncan McLay and Nick O’Brien from the Department of Primary Industries for their help with fish sampling. We also thank efishalbum.com for providing us with the fish images used in Figures 1 and 3, which were used with permission.

## Supplementary information

**SI Table 1.** Contig length and amino acid similarity of potentially novel vertebrate-associated viruses identified in this study.

**SI Table 2.** RNA sequencing library details, including sampling site, replicates and RNA concentration.

**SI Figure 1.** Phylogenetic relationships of the likely non-vertebrate viruses within the *Rhabdoviridae, Picornaviridae, Tombusviridae* and *Narnaviridae.* Viruses identified here are highlighted in blue. Maximum likelihood trees were estimated using amino acid sequences of the RdRp gene. Trees were mid-point rooted for clarity only and bootstrap values are represented as a percentage with branches scaled to amino acid substitutions per site. Tip labels represent virus name and NCBI/GenBank accession IDs.

**SI Figure 2.** Phylogenetic relationships of the likely non-vertebrate viruses within the *Nodaviridae, Phenuiviridae, Dicistroviridae* and *Permutotetraviridae.* Viruses identified here are highlighted in blue. Maximum likelihood trees were estimated using amino acid sequences of the RdRp gene. Trees were mid-point rooted for clarity only and bootstrap values are represented as a percentage with branches scaled to amino acid substitutions per site. Tip labels represent virus name and NCBI/GenBank accession IDs.

**SI Figure 3.** Phylogenetic relationships of bastrovirus sequences. Novel viruses are represented as black circles. Maximum likelihood tree was estimated using amino acid sequences of the ORF1 polypeptide. Trees were mid-point rooted for clarity only and bootstrap values are represented as a percentage with branches scaled to amino acid substitutions per site. Tip labels represent virus name and NCBI/GenBank accession.

