## Supplementary figures and images for "Metagenomic sequencing reveals a lack of virus exchange between native and invasive freshwater fish across the Murray-Darling Basin, Australia"

### SI Figure 1

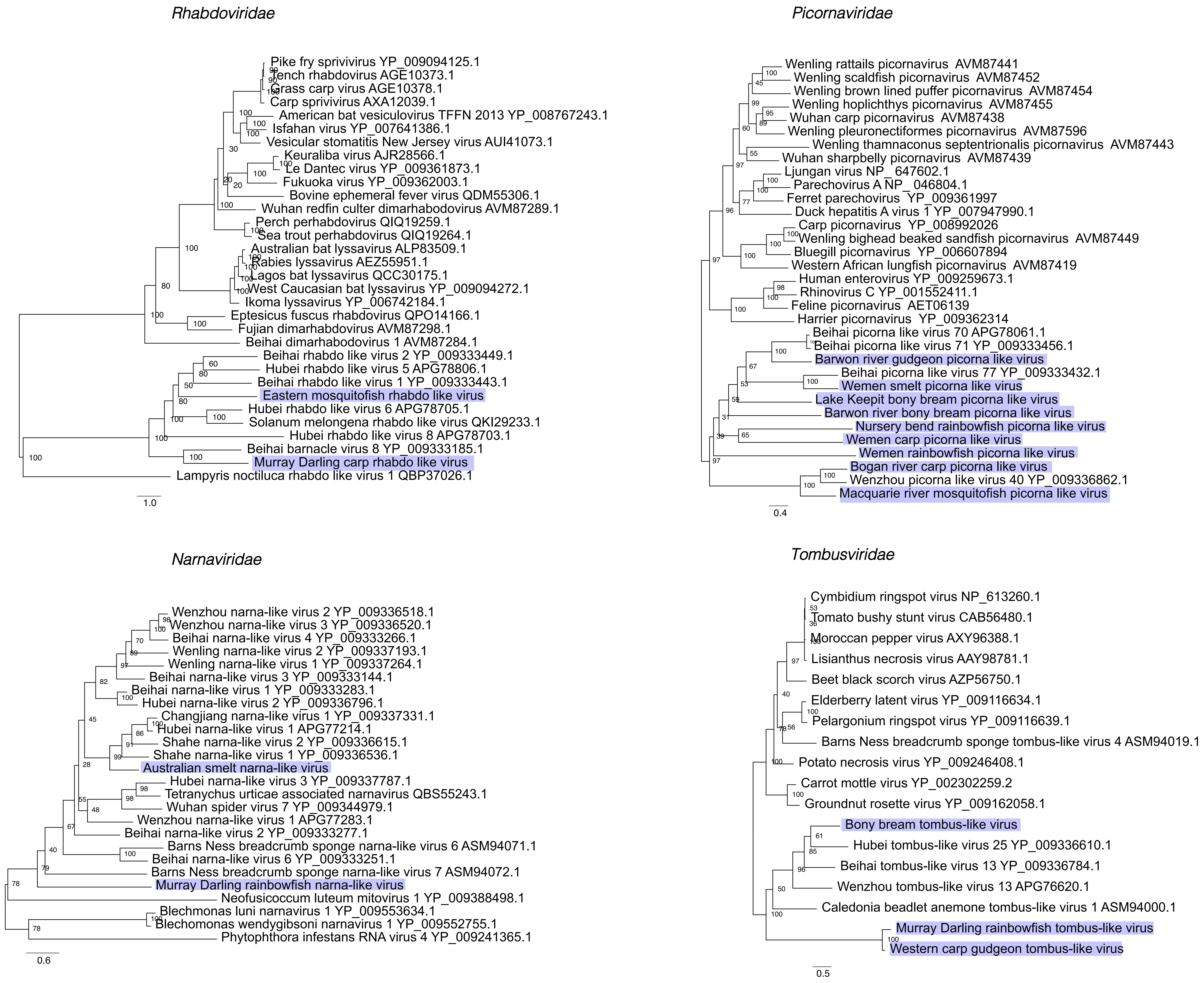

### SI Figure 2

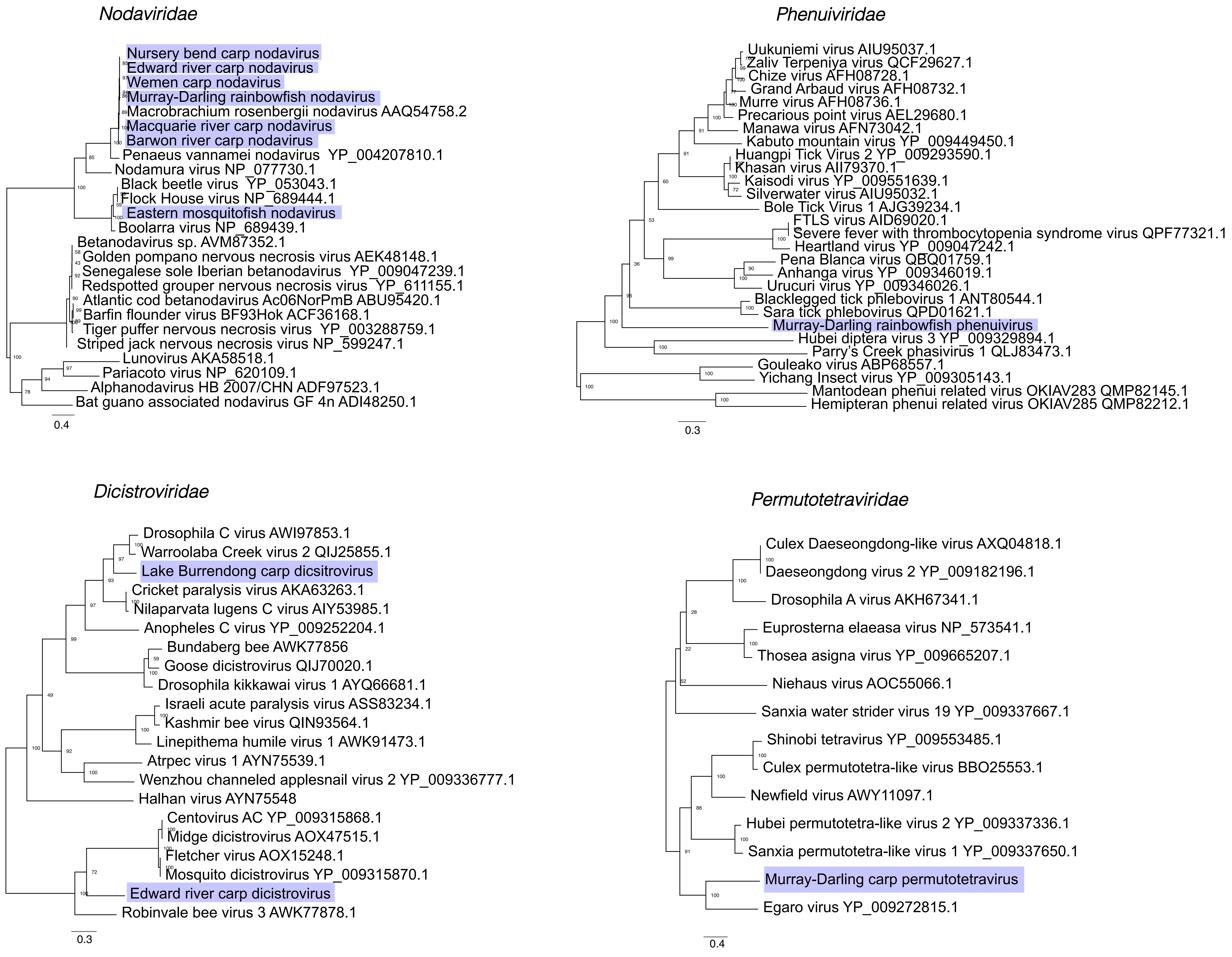

### SI Figure 3

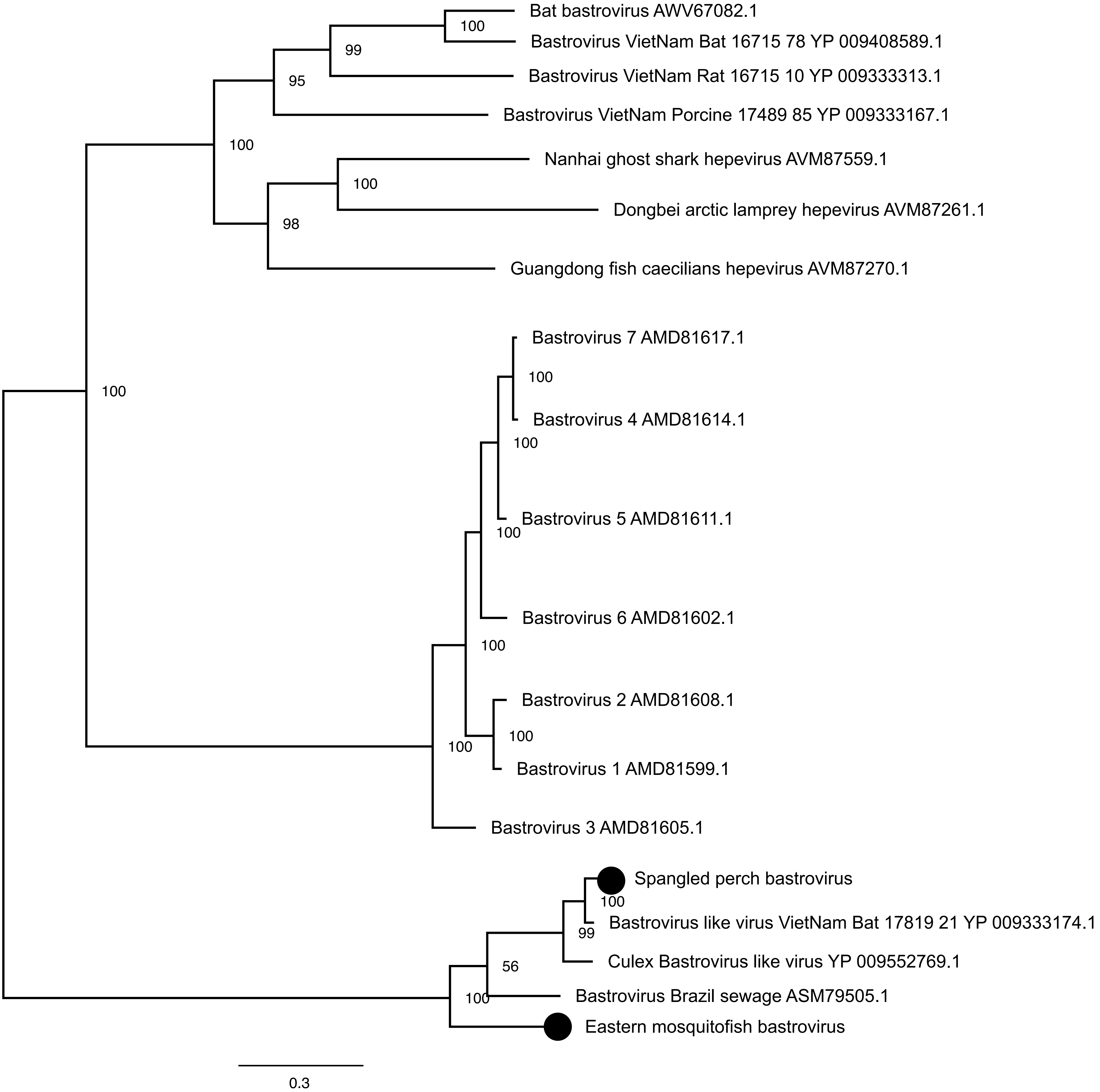
