## Supplementary material for "Metagenomic sequencing reveals a lack of virus exchange between native and invasive freshwater fish across the Murray-Darling Basin, Australia": SI Table 1

| Host | Virus name | Virus family | Gene/ORF | Length (nt) | Closest relative (NCBI/Genbank) | Amino acid similarity (%) |
|---|---|---|---|---|---|---|
| Western carp-gudgeon (*Hypseleotris spp.)* | *Western carp-gudgeon arenavirus* | *Arenaviridae* | L protein (RdRp) | 832 | *Wenling frogfish arenavirus 1* (YP_009551555) | 36.9 |
| Eastern mosquitofish (*Gambusia holbrooki*) | *Eastern mosquitofish arenavirus* | *Arenaviridae* | L protein (RdRp) | 5991 | *Wenling frogfish arenavirus 1* (YP_009551555) | 84.5 |
| | | | Nucleoprotein | 1884 | *Wenling frogfish arenavirus 1* (YP_009551555) | 78.7 |
| Spangled perch (*Leiopotherapon unicolor*) | *Spangled perch bastrovirus* | *Astroviridae* | Non-structural polyprotein | 1294 | *Bastrovirus-like virus Vietnam Bat* (YP_009333174.1) | 81.3 |
| Eastern mosquitofish (*Gambusia holbrooki*) | *Eastern mosquitofish bastrovirus* | *Astroviridae* | Non-structural polyprotein | 2463 | *Bastrovirus Brazil/sewage* (ASM79505) | 61.7 |
| | | | Structural polyprotein | 563 | *Bastrovirus Brazil/sewage* (ASM79506) | 75.4 |
| Murray-Darling rainbowfish (*Melanotaenia fluviatilis)* | *Murray-Darling rainbowfish astrovirus* | *Astroviridae* | RdRp | 1989 | *Wuhan astro-like virus* (AVM87125) | 40.3 |
| Common carp (*Cyprinus carpio*) | *Murray-Darling carp cultervirus* | *Bornaviridae* | L protein (RdRp) | 5190 | *Sharpbelly cultervirus* (AVM87541) | 93.3 |
| | | | Glycoprotein | 1575 | *Sharpbelly cultervirus* (AVM87539) | 86.8 |

| | | | | | | |
|---|---|---|---|---|---|---|
| | | | Nucleoprotein | 1095 | *Sharpbelly cultervirus* (AVM87536) | 92.9 |
| Bony herring (*Nematalosa erebi*) | *Bony herring calicivirus* | *Caliciviridae* | Polyprotein | 387 | *Atlantic salmon calicivirus* (AHX24377) | 80.3 |
| Common carp (*Cyprinus carpio*) | *Murray-Darling carp letovirus* | *Coronaviridae* | RdRp | 228 | *Pacific salmon nidovirus* (QEG08237) | 50.7 |
| Unspecked hardyhead (*Craterocephalus fulvus*) | *Hardyhead chuvirus* | *Chuviridae* | L protein (RdRp)<br><br>Glycoprotein<br><br>Nucleoprotein | 6363<br><br>1956<br><br>1566 | *Guangdong red-banded snake chuvirus-like virus* (AVM87272)<br><br>*Wenling fish chu-like virus* (AVM87276)<br><br>*Herr Frank virus* (QHX39758) | 44<br><br>41<br><br>34 |
| Western carp-gudgeon (*Hypseleotris spp.)* | *Western carp-gudgeon flavivirus* | *Flaviviridae* | NS5 | 1866 | *Cyclopterus lumpus virus* (ATQ64261) | 36 |
| Murray-Darling rainbowfish (*Melanotaenia fluviatilis*) | *Murray-Darling rainbowfish hantavirus* | *Hantaviridae* | RdRp | 1116 | *Bern perch virus* (QGM12349) | 27.3 |
| Common carp (*Cyprinus carpio*) | *Murray-Darling carp hepevirus* | *Hepeviridae* | Polyprotein (RdRp) | 2682 | *Cutthroat trout virus* (YP_004464929) | 31.1 |
| Eastern mosquitofish (*Gambusia holbrooki*) | *Eastern mosquitofish hepevirus* | *Hepeviridae* | Polyprotein (RdRp) | 6456 | *Wenling thamnaconus septentrionalis hepevirus* (AVM87557.1) | 29 |
| Western carp-gudgeon (*Hypseleotris spp.)* | *Western carp-gudgeon paramyxovirus* | *Paramyxoviridae* | RdRp | 512 | *Wenling tonguesole paramyxovirus* (AVM87378) | 35.2 |

| | | | | | | |
|---|---|---|---|---|---|---|
| Australian smelt (*Retropinna semoni*) | *Australian smelt picornavirus* | *Picornaviridae* | RdRp | 357 | *Eel picornavirus* (YP_008531322) | 54.2 |
| Western carp-gudgeon (*Hypseleotris spp.)* | *Western carp-gudgeon poxvirus* | *Poxviridae* | DNA polymerase | 213 | *Salmon gill poxvirus* (YP_009162448.1) | 61.4 |
| | | | DNA-dependant RNA polymerase subunit rpo22 | 340 | *Salmon gill poxvirus* (YP_009162433) | 46.5 |
| | | | DNA-dependant RNA polymerase subunit rpo19 | 226 | *Salmon gill poxvirus* (YP_009162475) | 40 |
| | | | DNA-binding virion core protein I1L | 643 | *Salmon gill poxvirus* (YP_009162452) | 28.1 |
| | | | Myristylated protein A16L | 247 | *Salmon gill poxvirus* (YP_009162490.1) | 32.9 |
| | | | Hypothetical protein (SGPV079) | 289 | *Salmon gill poxvirus* (YP_009162451.1) | 40.9 |
| Common carp (*Cyprinus carpio*) | *Murray-Darling carp reovirus* | *Reoviridae* | RdRp | 360 | *Wenling scaldfish reovirus (AVM87459)* | 40 |
| Common carp (*Cyprinus carpio)* | *Murray-Darling carp rhabdovirus* | *Rhabdoviridae* | RdRp | 592 | *Beihai dimarhabdovirus 1* (AVM87284) | 35.7 |
