## Supplementary material for "Metagenomic sequencing reveals a lack of virus exchange between native and invasive freshwater fish across the Murray-Darling Basin, Australia": SI Table 2

| Library | Site | Fish | Replicates | RNA concentration (ng/ul) |
|---|---|---|---|---|
| L1 | Lake Keepit | Common carp | 5 | 183.56 |
| L2 | Lake Keepit | Bony herring | 5 | 457.31 |
| L3 | Narrabri Creek | Common carp | 7 | 709.61 |
| L4 | Narrabri Creek | Bony herring | 4 | 537.85 |
| L5 | Narrabri Creek | Western carp-gudgeon | 5 | 451.44 |
| L6 | Narrabri Creek | Australian smelt | 5 | 153.54 |
| L7 | Narrabri Creek | Spangled perch | 1 | 339.90 |
| L8 | Narrabri Creek | Goldfish | 5 | 227.57 |
| L9 | Barwon River | Bony herring | 3 | 504.82 |
| L10 | Barwon River | Western carp-gudgeon | 4 | 378.86 |
| L11 | Barwon River | Common carp | 8 | 259.12 |
| L12 | Bogan River | Common carp | 8 | 427.19 |
| L13 | Bogan River | Bony herring | 5 | 929.30 |
| L14 | Bogan River | Western carp-gudgeon | 5 | 676.06 |
| L15 | Castlereagh River | Eastern mosquitofish | 9 | 173.52 |
| L16 | Castlereagh River | Western carp-gudgeon | 6 | 101.78 |
| L17 | Castlereagh River | Common carp | 10 | 716.04 |
| L18 | Macquarie River | Common carp | 10 | 202.85 |
| L19 | Macquarie River | Eastern mosquitofish | 6 | 355.86 |
| L20 | Gwydir River | Bony herring | 3 | 344.41 |
| L21 | Gwydir River | Common carp | 5 | 309.72 |

| | | | | |
|---|---|---|---|---|
| L22 | Lake Burrendong | Common carp | 5 | 137.24 |
| L23 | Abercrombie River | Common carp | 2 | 233.71 |
| L24 | Murray River (Nursery bend) | Murray-Darling rainbowfish | 5 | 113.42 |
| L25 | Murray River (Nursery bend) | Common carp | 4 | 167.15 |
| L26 | Edward River | Common carp | 4 | 519.99 |
| L27 | Edward River | Murray-Darling rainbowfish | 5 | 290.21 |
| L28 | Edward River | Australian smelt | 2 | 130.65 |
| L29 | Edward River | Unspecked hardyhead | 6 | 319.00 |
| L30 | Murray River (Wemen) | Murray-Darling rainbowfish | 3 | 148.63 |
| L31 | Murray River (Wemen) | Flat-headed gudgeon | 7 | 204.57 |
| L32 | Murray River (Wemen) | Australian smelt | 5 | 166.38 |
| L33 | Murray River (Wemen) | Common carp | 2 | 222.03 |
| L34 | Murray River (Coomealla) | Murray-Darling rainbowfish | 4 | 152.49 |
| L35 | Murray River (Coomealla) | Flat-headed gudgeon | 2 | 199.13 |
| L36 | Murray River (Coomealla) | Common carp | 4 | 179.50 |
